# An exon DNA element modulates heterochromatin spreading in the master regulator for sexual commitment in malaria parasites

**DOI:** 10.1101/2020.06.26.163105

**Authors:** Carlos Cordon-Obras, Anna Barcons-Simon, Christine Scheidig-Benatar, Aurelie Claës, Valentin Sabatet, Damarys Loew, Artur Scherf

## Abstract

Heterochromatin is essential in all eukaryotes to maintain genome integrity, long-term gene repression and to help chromosome segregation during mitosis. However, heterochromatin regions must be restricted by boundary elements to avoid its spreading over actively transcribed loci. In *Plasmodium falciparum*, facultative heterochromatin is important to regulate parasite virulence, antigenic variation and transmission. However, the underlying molecular mechanisms regulating repressive regions remain unknown. To investigate this topic, we chose the *ap2-g* gene, which forms a strictly delimited and independent heterochromatin island. Using electrophoretic motility shift assay (EMSA) we identified an *ap2-g* exon element at the 3’ end binding nuclear protein complexes. Upon replacement of this region by a *gfp* gene, we observed a shift in the heterochromatin boundary resulting in HP1 (Heterochromatin Protein 1) spreading over ∼2 additional kb downstream. We used this DNA element to purify candidate proteins followed by proteomic analysis. The identified complexes were found to be enriched in RNA-binding proteins, pointing to a potential role of RNA in the regulation of the *ap2-g* 3’ heterochromatin boundary. Our results provide insight into the unexplored topic of heterochromatin biology in *P. falciparum* and identify a DNA element within the master regulator of sexual commitment modulating heterochromatin spreading.

## INTRODUCTION

The apicomplexan parasite *Plasmodium falciparum* is the causative agent of the deadliest and more prevalent form of human malaria. This disease still stands as one of the major health constraints for development of several endemic areas, infecting more than 200 million people and causing almost a half million deaths per year, especially in Africa^1^. During its complex life cycle, the parasite undergoes dramatic transcriptional changes in response to the different environments it must face in its arthropod vector, mosquitoes from the genus *Anopheles*, and human host^2–4^. In the latter, *P. falciparum* first infects hepatocytes before progressing to red blood cells (RBCs) where chronic infection occurs. There, the parasites undergo a 48-hour life cycle whereby they multiply and release 12-30 daughter cells (merozoites) in the bloodstream able to re-invade new RBCs.

Life-cycle specific gene expression is controlled at the transcriptional level^5^ and modulated by various epigenetic layers^6–8^. In addition, various types of post-transcriptional control mechanisms emerge as important regulators of protein expression^9^.

A particular histone post-translational modification, H3K9me3, attracted the attention of researchers early on being that it is the landing pad for Heterochromatin Protein 1 (PfHP1), ortholog of *Schizosaccharomyces pombe* Swi6^10^, and essential for heterochromatin formation. This chromatin state, characterised for dense nucleosome packing and refractoriness to transcription, seems to play a pivotal role in gene expression regulation of parasite-host crosstalk. Globally, around 400 genes are enriched with H3K9me3/HP1 in *P. falciparum*. The vast majority are included in clusters whose boundaries mark synteny breakpoints or species-specific indels, and most of them belong to clonally variant gene (CVG) families^11,12^. Among them, the best studied are the *var* genes, which encode the PfEMP1 surface antigens, one of the major virulence factors involved in cytoadhesion and pathogenesis^13^. A notable exception to the aforementioned generalities is AP2-G, a transcription factor (TF) proven to be a master regulator for sexual commitment in *Plasmodium* genus^2,14,15,16,17^. The *ap2-g* gene remains repressed in most blood stage parasites during the intraerythrocytic developmental cycle (IDC), by means of heterochromatinization, and is only activated in a small subset of cells by specific removal of HP1 via GDV1^18^. The *ap2-g* locus does not belong to any heterochromatic multigene cluster but forms an isolated heterochromatin island in the central region of chromosome 12. It appears to be the only TF repressed by heterochromatin, a phenotype conserved in all *Plasmodium* species analysed so far^19^. This makes the *ap2-g* gene an appealing model for studying the biology of heterochromatin in *P. falciparum*, a largely unexplored field.

Aside from HP1 distribution across the genome, we have very few clues about heterochromatin assembly in the parasite and the elements that avoid its undesired spreading. Such DNA sequences are usually recognised in a specific manner by factors that counteract the required chromatin environment for heterochromatin propagation, or impose physical limitations to avoid it^20^. Boundary elements have been suggested in *P. falciparum*^11,12^ but never experimentally validated. In *ap2-g* heterochromatin is restricted to the 5’ UTR, the ORF and sharply drops in the 3’ end of the gene, suggesting a precise barrier effect in this region.

Here, we identified a DNA element at the 3’ terminal end of *ap2-g* that associates with nuclear proteins in gel shift experiments. Replacement of the 3’ region exon region by a *gfp* gene, shifted the heterochromatin boundary by ∼2 kb downstream. Candidate regulatory proteins that bind to this element were identified by affinity purification and mass spectrometry analysis. We observed an enrichment for RNA-binding proteins pointing to a potential association of RNA to this DNA element.

## RESULTS

### A protein complex specifically binds to the 3’ end of *ap2-g* gene

In order to determine the factors potentially involved in the boundary at the 3’ end of *ap2-g* gene, we performed a screening, searching for proteins able to bind to that region in a sequence-specific manner. To do so, we used biotinylated overlapping DNA fragments of at least 20 bp as probes for EMSA. Targeted regions spanned from -765 to +464 bp, respective to the stop codon. All primers employed in this work are listed in Table S1.

Our analysis revealed only one fragment yielding a shifted band and responding to competition with a non-biotinylated probe (100-fold excess). This fragment located inside the ORF of *ap2-g*, downstream the AP2 domain (Fig. 1a). A shorter fragment (EMSA 2.2) still kept the shifted band (183 bp), indicating a protein complex interacting with this DNA element (Fig. 1b). We referred to it as ap2-g-3’PE (ap2-g 3’protein-recruiting element). However, our attempts to achieve shorter fragments keeping the complex binding capacity were not successful (Fig. 1b).

**Figure 1.**
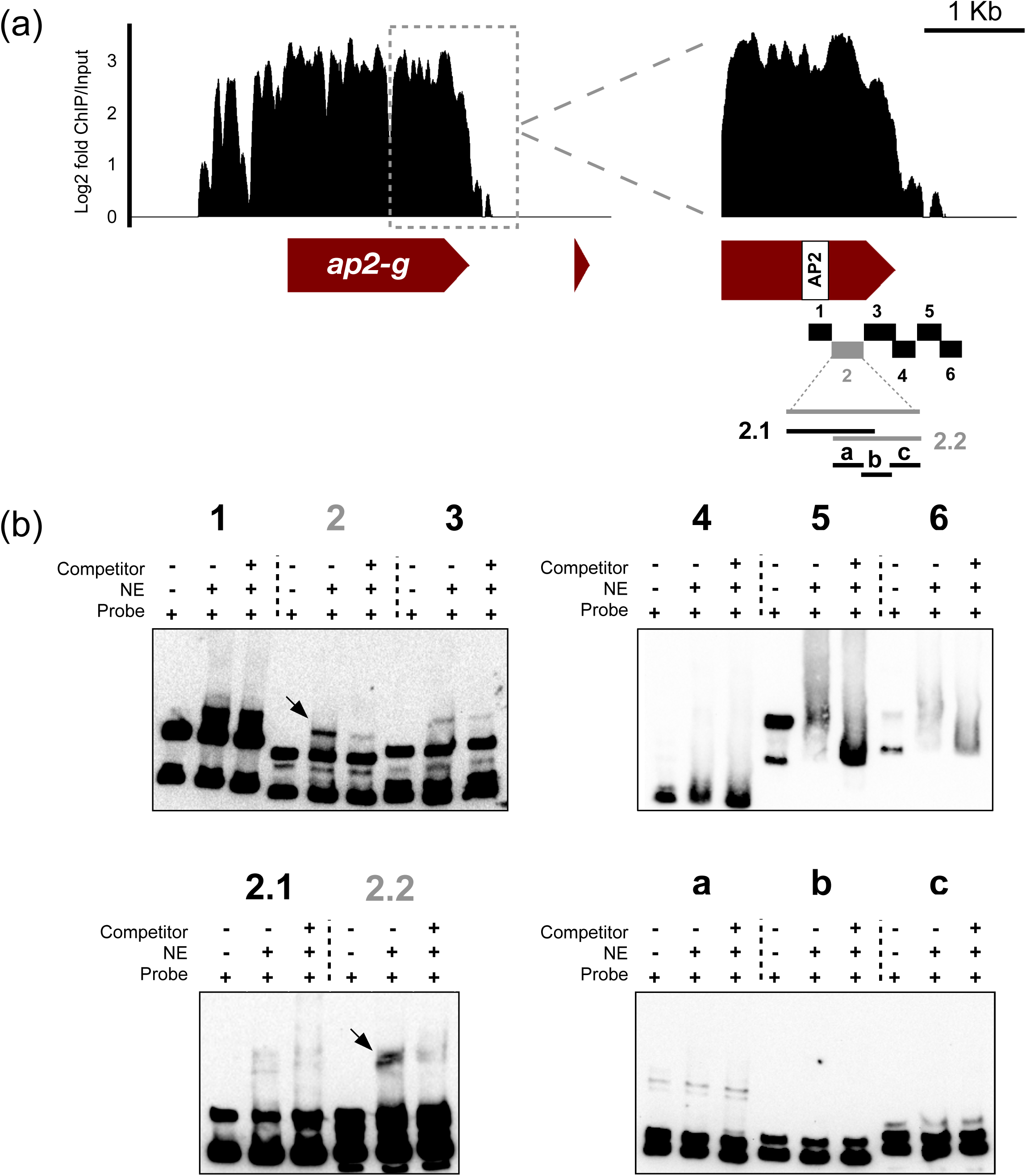
Search of protein complex binding to the 3’ end of *ap2-g* locus. (a) Left: *ap2-g* ORF and HP1 ChIP-Seq signal in log2 fold increased over input. Right: zoomed view of the 3’ end of *ap2-g* ORF and scheme depicting the fragments tested by EMSA. Sequences in grey mark a positive shift. (b) EMSA of all fragments. Arrows point the specific shift bands which fade upon competition.

### Replacement of the *ap2-g* 3’ end extends the heterochromatin boundary

In order to gain insight into the function of the DNA element found to bind a specific protein complex, we aimed to engineer a line in which such fragment was depleted. However, given that nucleation centres for heterochromatin assembly are unknown, we anticipated a potential problem whereby genome edited parasites could eventually differentiate into gametocytes if we interfere with chromatin structure of *ap2-g*. In order, to avoid this problem, we replaced the last 765 bp of *ap2-g* with a *gfp* gene, including the AP2 domain. In doing so, this would mimic the natural mutation of GNP-A4 clone, known to be unable to accomplish gametocytogenesis^15^. Transfection of NF54 A11 clone with pL6-gfp-ap2g and pUF-Cas9 plasmids resulted in such a line (AP2-G_gfp_ line hereon) (Fig. 2a). The correct insertion was verified by PCR using specific primers targeting *gfp, ap2-g* and 3’UTR (Fig. 2b).

**Figure 2.**
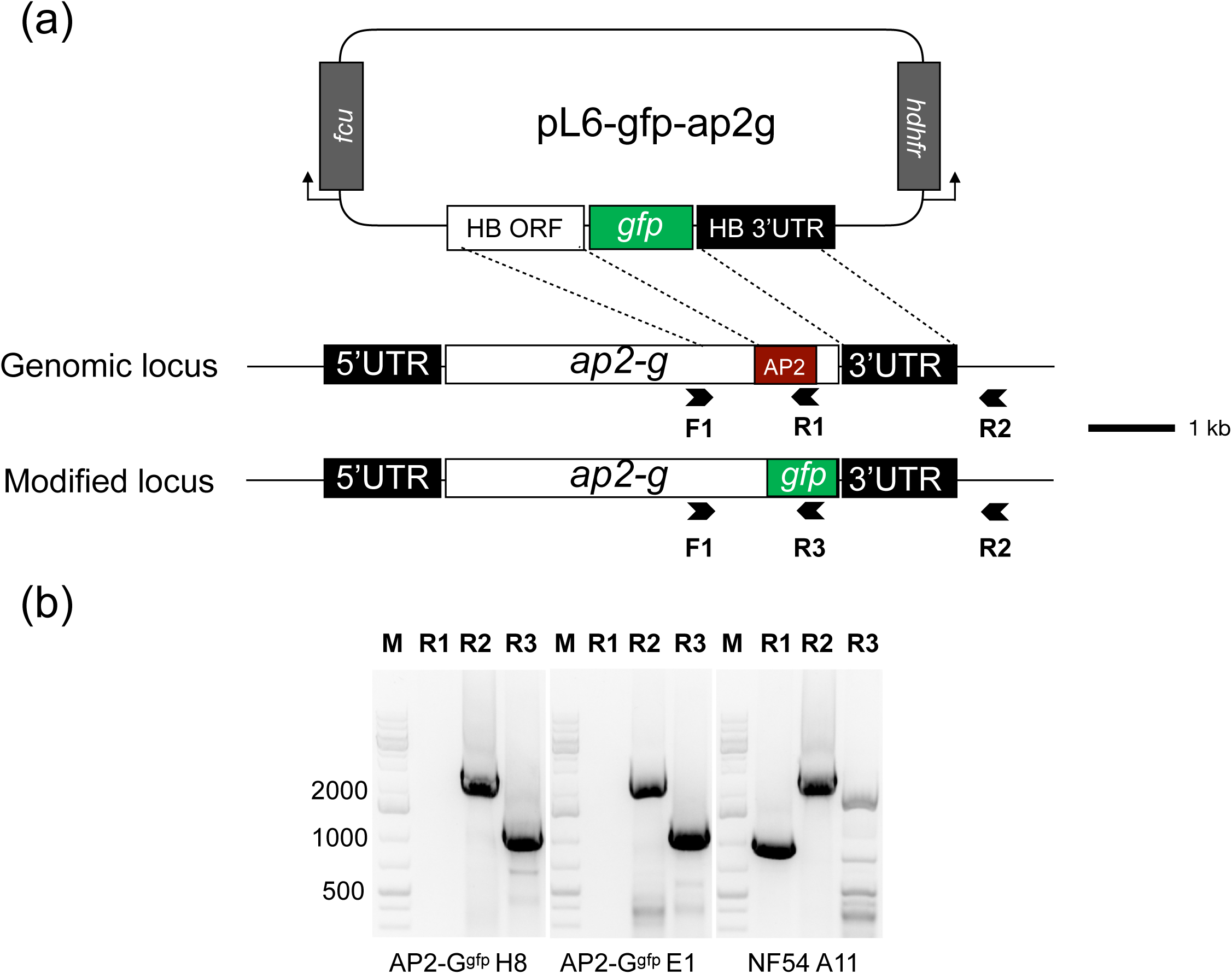
Development and validation of AP2-G_gfp_ line. (a) Depiction of the pL6-gfp-ap2g plasmid co-transfected with pUF-Cas9 to replace the last 765 bp of the *ap2-g* ORF by *gfp* gene. Arrows mark the primers used for further validation. Red square marks the AP2 domain, included in the replaced region. HB: homology box. (b) PCR validation of correct integration for AP2-G_gfp_ clones and parental line. Forward primer was F1 in all PCRs.

We further checked the phenotype of two clones of the AP2-G_gfp_ line. Consistent with predictions, upon induction, clones H8 and E1 yielded no gametocytes, whereas the parental line reached ∼8% conversion rate (Fig. 3a and 3b). GFP was visible on live imaged cells at a very low rate (<0.1%), and we were never able to obtain higher ratios of expressers even after gametocyte induction (Fig. 3c), indicating that the deleted 3’ exon region contributes to the expression of AP2-G. Furthermore, transcription of *ap2-g* varied in AP2-G_gfp_ clones, peaking in late ring stages rather than in early rings and schizonts as occurred in the parental line (Fig. 3d) as previously observed in other clones unable to sexual differentiation^14^. Next, we tried to sort the GFP-expressing cells by flow cytometry in order to further explore the *ap2-g* activated parasite development and transcriptome. However, the GFP signal appeared too low to be unambiguously detected with this technique and we were unable to distinguish GFP expressers from autofluorescence cells present in the wild type strain (Fig. S1).

**Figure 3.**
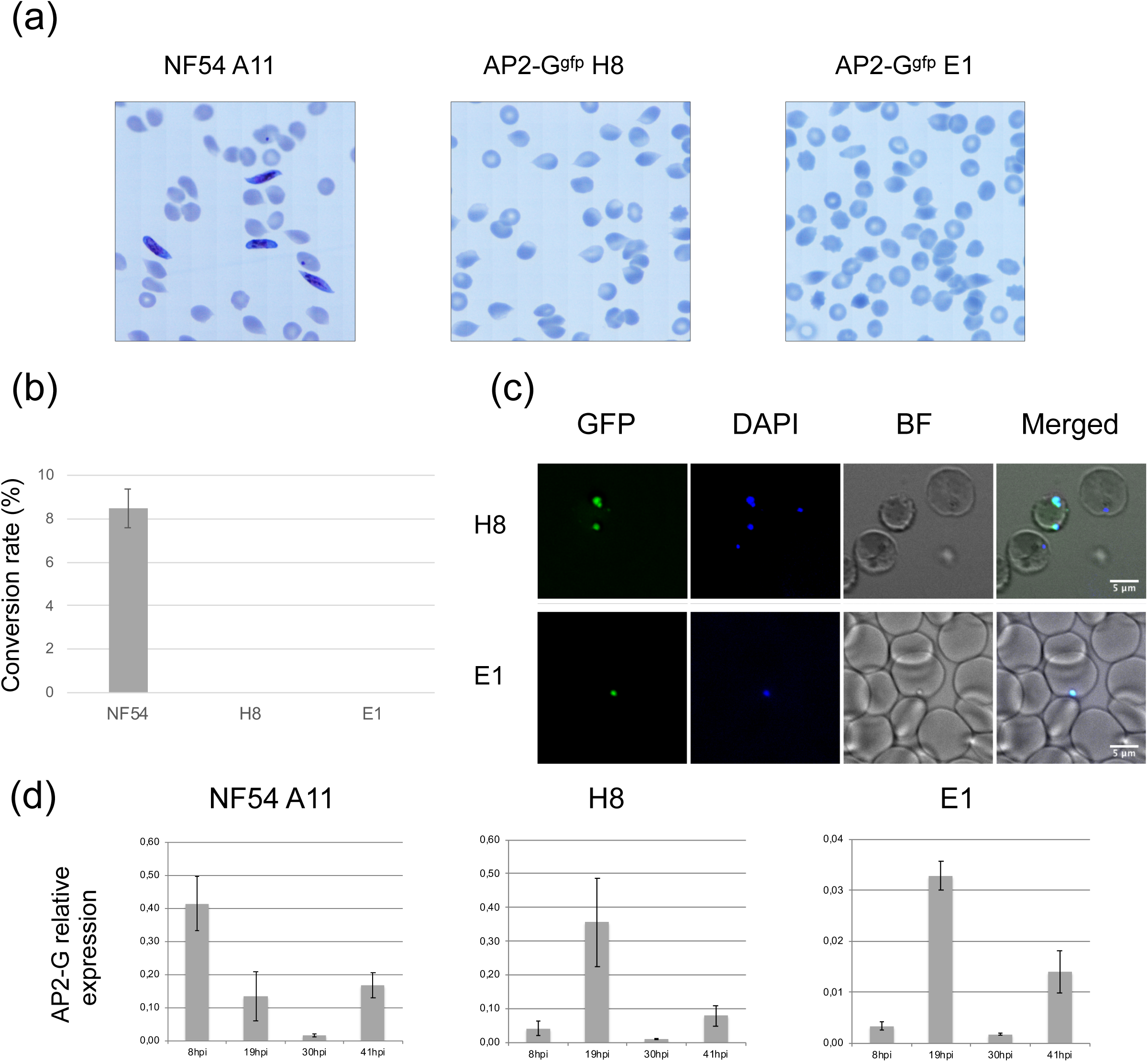
AP2-Ggfp line phenotype validation. (a) Giemsa staining of induced gametocytes of parental line NF54 A11 and two clones of AP2-Ggfp line. (b) Gametocyte conversion rate of lines in (a). Mean ± SEM of three independent experiments is shown. (c) Live imaging of AP2-Ggfp clones. Scale bar at 5 µm. (d) Expression levels of AP2-G at different time points measured by RT-qPCR. Transcript levels are shown relative to ubiquitin-conjugating enzyme (PF3D7_0812600). Mean ± SEM of two independent experiments is shown.

To evaluate the impact of replacing the terminal part of *ap2-g* on the heterochromatin profile, we performed ChIP-Seq with anti-HP1 on tightly synchronised 36 hours post invasion (hpi) parasites from both parental and two clones of the AP2-G_gfp_ lines. In NF54 A11, HP1 accumulation extends over the complete ORF, dropping sharply after the stop codon. Accumulation extends ∼3 kb upstream of the start codon, progressively falling after a first drop and increases again at ∼1.7-1.9 kb from the start codon. AP2-G_gfp_ clones exhibited a similar pattern upstream the *ap2-g* ORF, except the HP1 accumulation extends after the stop codon ∼2 kb beyond the limit shown in the parental line (Fig. 4a). Interestingly, available ATAC-Seq data^21^ revealed accessible chromatin where HP1 accumulation stops in the AP2-G_gfp_ clones, pointing to a chromatin environment that may halt HP1 spreading further downstream of *ap2-g*. However, in the immediate downstream vicinity the *ap2-g* ORF does not show an open chromatin region (Fig. 4a), supporting a direct role of the identified 3’ exon region in restricting spreading of HP1.

**Figure 4.**
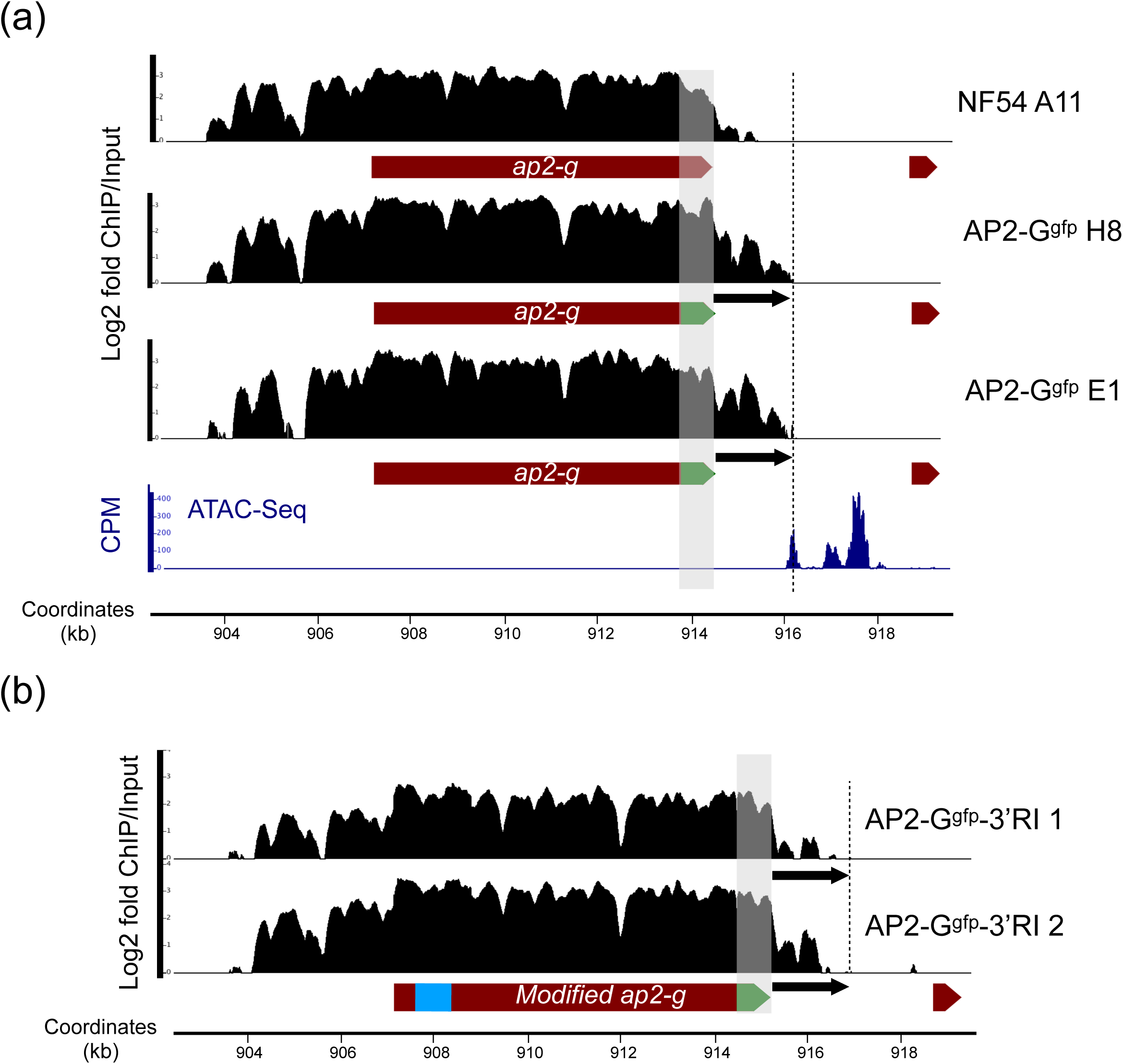
Heterochromatin boundary shift in AP2-Ggfp clones. (a) ChIP-Seq with anti-HP1 antibodies expressed as log2 fold enrichment over input. ATAC-Seq data from Ruiz et al.21 is shown in dark blue as normalised tags (counts per million reads -CPM-). Region replaced in AP2-Ggfp lines is shaded in grey. One replicate experiment per each clone is shown. (b) Same as (a) but using the line AP2-Ggfp-3’RI. Results for two clones are shown. Re-inserted 3’ end is marked in blue. Distance between stop codon and shifted heterochromatin boundary in AP2-Ggfp clones is shown as an arrow and a vertical dashed line in both (a) and (b) for comparison. Gfp gene is marked in green.

### Insertion of *ap2-g* 3’ end in AP2-G_gfp_ restricts the 3’ heterochromatin boundary

In order to explore the role as boundary of the elements contained in *ap2-g* 3’ end, we re-inserted it in the 5’ end of the modified *ap2-g-gfp* locus (AP2-G_gfp_-3’RI line). To do so, we transfected the AP2-G_gfp_ E1 clone with pL6-ap2g-3’RI and pUF-Cas9 plasmid as described in Material and Methods section. We predicted that the heterochromatin spreading would interrupted at the insertion. Strikingly, the re-insertion of *ap2-g* 3’ end did not disrupt the HP1 profile in the exon but, restored the boundary at the 3’ end of the ORF with a profile similar to the wild type NF54 A11 in two independent experiments (Fig. 4b). We conclude the ap2-g-3’PE can act over a certain distance and is not restricted to a precise location.

### Identification of proteins bound to ap2-g-3’PE by mass spectrometry analysis

In order to determine the potential factors bound to the ap2-g-3’PE, we used the biotin probes that yielded positive EMSA results (see Fig. 1) to purify their interacting proteins. For that, we employed the ap2-g-3’PE (2.2) and the 2.1 as control. Quantitative label-free mass spectrometry analysis (ap2-g-3’PE versus control) yielded fifty-eight proteins enriched with a two-fold threshold or unique to the ap2-g-3’PE fragment.

Among the significant interactors, we found seventeen nucleic acid binding proteins (GO:0003676) and seven uncharacterised proteins, based on Gene Ontology annotation terms (Fig. 5 and Table S2). Neither enriched nor unique quantified peptides were found in the control fragment. These results raise the possibility that a complex set of ap2-g-3’PE-interactors is involved in the observed phenotype of the *ap2-g* gene heterochromatin remodelling and regulation of induction of sexual commitment.

**Figure 5.**
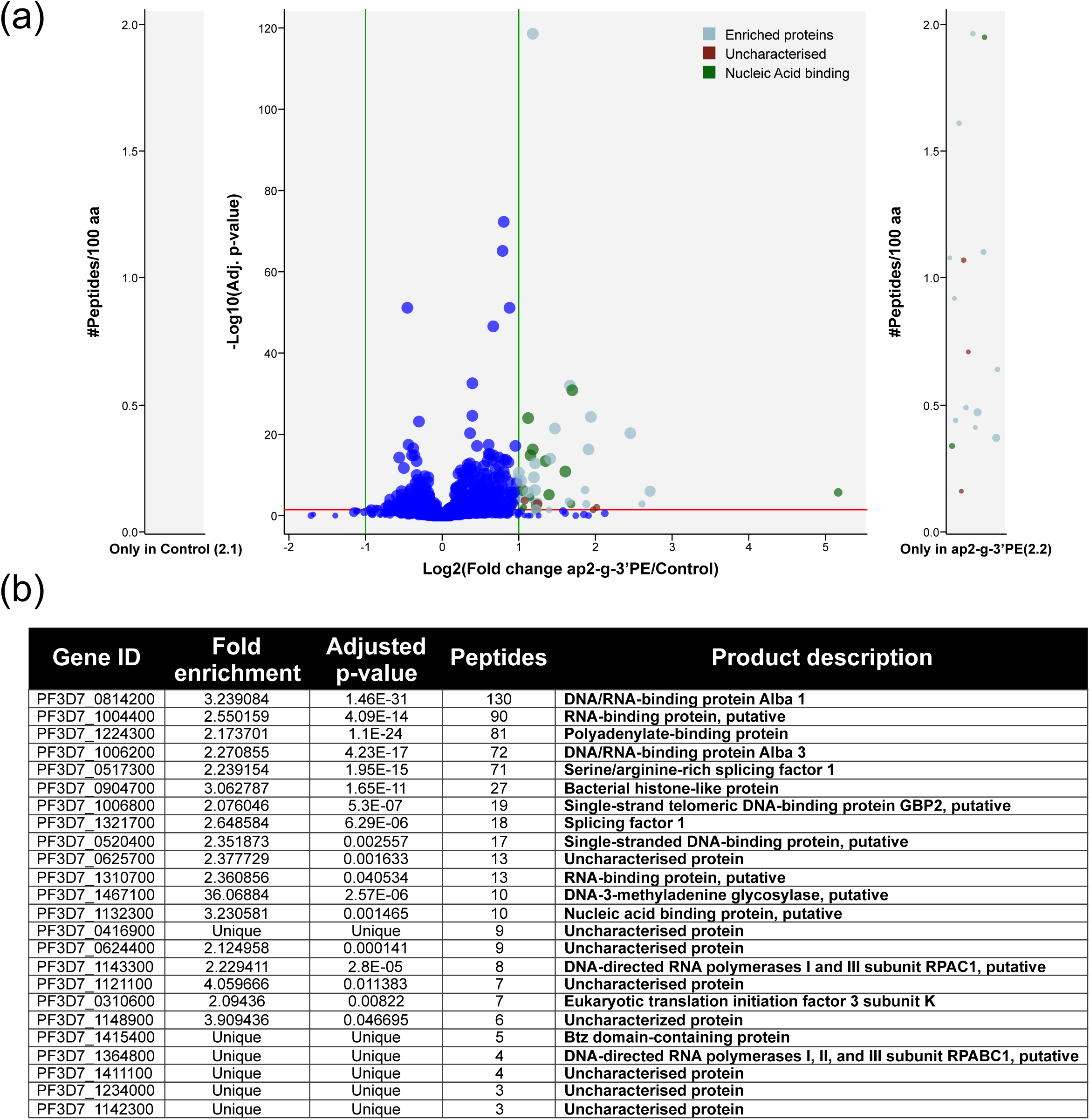
Proteomics mass spectrometry identification of factors enriched in terminal 3’ region of AP2-G. (a) Volcano plot showing differential enrichment of proteins bound to 2.2 (ap2-g-3’PE) and 2.1 (control) fragments obtained by using quantitative label-free mass spectrometry analysis performed from five replicates. Shown are the fold changes (sample vs control), quantified with an absolute fold change of 2 or more with an adjusted p-value of ratio significance of 0.05 or less and with 3 or more peptides. Nucleic acid binding proteins are highlighted in green along with uncharacterised ones (red). Proteins only found in sample are shown at the right panel. (b) List of nucleic acid binding proteins and uncharacterised proteins found in the quantitative label-free mass spectrometry analysis.

## DISCUSSION

Facultative heterochromatin is a major regulator of gene expression during the life cycle of malaria parasites. It is of particular importance to regulate important pathogen processes that depend on phenotypic plasticity such as immune evasion and transmission. Heterochromatin silenced gene families occupy sub-telomeric regions as well as some internal chromosome clusters. Heterochromatin distribution varies during life cycle and between different stages. For instance, HP1 expands further into sub-telomeric coding regions in sporozoites, leading to silencing of asexual blood stage genes that are not expressed in sporozoites ^19,22^. The molecular events that control boundaries in this protozoan pathogen, however, remain elusive. In this work, we used the *ap2-g* gene as a model to explore factors that set the limits of heterochromatin. HP1 forms a well delineated island around this master regulator of sexual commitment, located in a central euchromatin chromosome region. At each asexual blood stage cycle, a small subset of parasites de-represses the *ap2-g* gene, leading to the activation of a transcriptional cascade that initiates gametocytogenesis. Noteworthy, identified DNA motif by *P. falciparum* AP2-G is also highly conserved in the *P. berghei* ortholog ^17^ and other apicomplexa as *Theileria annulata* 23, indicating a similar biological role. However, no significant homology was detected in other genes that are under control of heterochromatin in the silent state.

HP1 occupation sharply drops at the 3’ end of *ap2-g*, suggesting the presence of a boundary element restricting its spreading. In order to determine the regions involved in this putative barrier activity we performed EMSA experiments using specific overlapping probes surrounding the 3’ end of *ap2-g*. Our results showed a specific DNA-protein interaction with a 183 bp probe downstream of the AP2 DNA-binding domain, called ap2-g-3’PE (Fig. 1). We aimed to explore the effective involvement of this region in the HP1 distribution by replacing the 3’ end (765 bp of exon of the *ap2-g* gene) with the *gfp* gene. We reasoned that depletion of short sequences leaving the AP2 domain intact could potentially de-repress the *ap2-g* and lead to gametocyte differentiation making recovery of transfectants not feasible in *P. falciparum*. Our engineered line, AP2-G_gfp_, displayed the expected phenotype of loss of functional AP2-G, rendering parasites unable to trigger gametocytogenesis and with a noticeable change in the transcriptional pattern across the IDC (Fig. 3). Similar features were observed previously with the clone F12^14^, a natural mutant for the *ap2-g* that has lost its ability to differentiate to gametocytes. We detected GFP expresser cells in live imaging, confirming that a subset of parasites was able to initiate the transcriptional cascade for differentiation, but our attempts to sort them were unsuccessful due to the low number and intensity of the signal (Fig. S1). ChIP-Seq analysis on the AP2-G_gfp_ line revealed a heterochromatin boundary shift in the 3’ end upon replacement of the last 765 bp of *ap2-g* ORF by *gfp*. HP1 distribution extended ∼2 additional kb downstream respective to the profile displayed by the parental line (Fig. 4a). Interestingly, in both AP2-G_gfp_ clones, the heterochromatin domain ends at an open chromatin region, as proven by available ATAC-Seq data^21^, likely containing the promoter of the nearby gene (PF3D7_1222700). Such regions are known to hinder the HP1 spreading, as they tend to lack nucleosomes, negating the histone mark required for HP1 binding^24^. There is no evidence of open chromatin in the immediate vicinity of *ap2-g* end and, therefore, the sharp drop of HP1 in the parental line cannot be attributed to this mechanism.

In order to confirm any possible boundary element within *ap2-g* 3’ end, we re-inserted it in the 5’ end of the modified *ap2-g-gfp* locus (>5 kp upstream of the original location). To our surprise, we observed no disruption of heterochromatin organisation at the point of insertion. However, the 3’ HP1 occupation was almost entirely restored to the *ap2-g* wild type (Fig 4b). The fact that the DNA element influenced the heterochromatin distribution at a distance > 5kb point to a long-range effect, rather than a strict location dependency as usually found in eukaryotes^20^. Noteworthy, it is not common that a chromatin boundary element is found within an ORF in most eukaryotic systems. However, it has been previously speculated that chromatin regulatory elements may exist within exons in *P. falciparum*. It was reported that *eba-140* gene is divided in two different chromatin domains, the one from the 5’ appears to control the *eba-140* transcription, whereas the one in the 3’ terminal shares the chromatin state with the neighbouring gene^25^. No further analyses were conducted to verify this hypothesis. However, this points to the existence of DNA elements within coding sequences that regulate chromatin states beyond the *ap2-g* gene.

In order to elucidate the factors involved in the boundary function found in the *ap2-g* gene, we purified the protein complex bound to ap2-g-3’PE region and performed proteomics and quantitative label-free mass spectrometry analysis. We did not observe any enrichment of proteins such as Sir2 that have been previously shown to specifically unlock heterochromatin at a particular *var* gene loci^26,27^. However, we found two Alba members, able to bind DNA and RNA^28,29^, and several other RNA-binding proteins of unknown function. None of them have been involved in chromatin remodelling so far. At this stage we can only speculate that the enrichment of these factors may point to the association of RNA in the complex that forms onto *ap2-g-3’PE*. Validation experiments of those candidates are needed to confirm their recruitment to the *ap2-g* locus and explore their role (including RNA) in the observed modulation of HP1 occupation. In vivo isolation of a specific chromatin region has been reported recently using dead Cas9^30^ and this technique may help further explore the native protein/RNA complex that interacts with ap2-g-3’PE.

In conclusion, our work has identified a DNA element in the *ap2-g* exon 3’ region that recruits a protein complex enriched for RNA-binding proteins. Replacement of the ap2-g-3’PE element by a *gfp* sequence induces an extension of the heterochromatin boundary by additional 2 kb. Insertion of ap2-g-3’PE 5’ upstream of the original location in the mutated *ap2-g* exon reverses the HP1 occupation to wild-type like situation. These results strongly point to the existence of exon encoded regulatory elements of chromatin state in malaria parasites and provide a first insight into the mechanism of heterochromatin spreading in this pathogen.

## METHODS

### Parasites culture and synchronisation

Blood stage NF54 A11 clone *P. falciparum* parasites were cultured in human O+ red blood cells in RPMI-1640 medium supplemented with Albumax II (10% v/v), hypoxanthine (200 μM) and gentamycin (50 μg/mL) in 3% CO2 and 5% O_2_ at 37 °C. Human blood was obtained from informed donors in the EFS (Établissement Français du Sang -French Blood Establishment-) under authorization number HS 2016-24803. Parasites were synchronised with a 6h time window by 5% sorbitol lysis (Sigma) during ring stage, followed by plasmagel enrichment in schizont stage and another sorbitol treatment 6 h after. Synchronized parasites were harvested at 3.3% haematocrit and ∼2-4% parasitaemia. Parasite development was monitored by Giemsa staining.

### Plasmids, transfections and transgenic lines

The plasmid used for replacement of *ap2-g* 3’ end and insertion of ap2-g 3’end in 5’ of *ap2-g* derived from previously described pL6-GFP^31^. Homology boxes, obtained through sequential PCRs were inserted into the pL6 vector digested at SacII and AfIII sites. The guide for Cas9 was obtained by direct hybridization of the oligos and inserted in the same plasmid by Gibson assembly (In-Fusion HD cloning kit; Clontech) using the BtgZI sites.

The pL6 derived plasmids and pUF-Cas9^31^ were co-transfected (25 μg each) in the NF54 A11 clone (for 3’ replacement) or AP2-G_gfp_ E1 clone (for 5’ insertion). To do so, ring stage parasites were transfected following the protocol described elsewhere^32^ and maintained under WR99210 and DSM1 drug selection pressure. Parasite clones were obtained by limiting dilution.

### Gametocyte conversion assay and live cell imaging

Gametocytogenesis was performed as previously described^33^. For live cell imaging, the bottom of imaging dishes (Ibidi) were covered with concanavalin A (5mg/mL) and incubated at 37°C for 30 min. Next, concanavalin A was carefully removed and dishes gently were washed with sterile PBS. 500 μL of parasite cultures (3-5% parasitaemia, 3-4% haematocrit), were then added to the dish and let to settle for 10 minutes at 37°C. Unattached cells were washed out with PBS and finally covered with culture medium prepared with phenol red free RPMI 1640. Samples were visualized using a Deltavision Elite imaging system (GE Healthcare) and processed with the Fiji package (http://fiji.sc).

### RNA isolation and reverse transcription quantitative PCR (RT-qPCR)

RNA was harvested as previously described^34^ from synchronised parasite cultures at indicated timepoints after lysis with 0.075% saponin in PBS followed by one wash in PBS and resuspension in Qiazol. Total RNA was extracted using the miRNeasy mini kit, performing on-column DNase treatment (Qiagen). Reverse transcription was achieved using SuperScript VILO (Thermo Fisher Scientific) and random hexamer primers. cDNA levels were measured by quantitative PCR in the CFX384 real time PCR detection system (BioRad) using Power SYBR Green PCR Master Mix (Applied Biosystems) and specific primers^34^. Starting quantity means of three replicates were extrapolated from a standard curve of serial dilutions of genomic DNA and normalised with a housekeeping gene (ubiquitin-conjugating enzyme, PF3D7_0812600).

### Chromatin immunoprecipitation and data analysis

ChIP was performed as previously described^22^ with parasites synchronised at 36 hpi. Sonicated chromatin (500 ng DNA content) was immunoprecipitated overnight with 0.5 μg of anti-HP1 (Genscript) polyclonal rabbit antibody^22^ and incubated after with 25 μL of Dynabeads Protein G (Invitrogen) for two hours. Subsequent washing, reverse cross-linking and DNA extraction were carried out as described before^22^. Sequencing libraries were prepared with the immunoprecipitated DNA using the MicroPlex Library Preparation Kit v2 (Diagenode) with the KAPA HiFi polymerase (Kapa Biosystems) for the PCR amplification. For each ChIP sample, a control DNA corresponding to the ChIP input was processed in parallel. Multiplexed libraries were subjected to 150 bp paired-end sequencing on a NextSeq 500 (Illumina)^34^. Fastq files were obtained by demultiplexing the data using bcl2fastq (Illumina) prior to downstream analysis. A minimum of two biological replicates were analysed for each clone.

Sequencing reads were mapped to the *P. falciparum* genome^35^(PlasmoDB, v29) using Burrows-Wheeler Alignment tool (BWA-MEM) with default settings^36^. PCR duplicates were removed. ChIP-seq data was normalised over input and peak calling was performed using the MACS2^37^ software with default parameters and a 0.05 false discovery rate (FDR) cut-off.

### Protein nuclear extract, probes and electrophoretic motility shift assay (EMSA)

Nuclei were obtained from NF54 A11 parasites treated with 0.15% saponin and incubated in rotation for 1 hour at 4° C with 1x low salt lysis buffer (20 mM Tris HCl pH 7.5, 10 mM KCl, 2 mM DTT, 1.5 mM MgCl_2_, 1% Triton X-100 and 1x proteinase inhibitor cocktail -PIC-). After spinning at 17,000 g for 20 min at 4° C, nuclei were extracted with high salt 1x lysis buffer (20 mM Tris HCl pH 7.5, 600 mM KCl, 2 mM DTT, 1.5 mM MgCl_2_, 1% Triton X-100 and 1x PIC) for 30 min in rotation at 4° C and brief sonication. Samples were spun again and supernatant was taken and diluted four times in low salt lysis buffer without detergent. Protein concentration was measured by Bradford assay, performing a reference curve by increasing BSA concentrations.

Probes for EMSA were obtained by direct PCR amplification of desired fragments with biotin labelled oligos (5’ biotin modification in forward primer), from genomic DNA of NF54 A11 parasites. PCR products were separated in a 2% agarose gel, sliced and purified with Nucleospin PCR and Gel Clean-up Kit (Macherey-Nagel). Concentration was measured in a NanoDrop device.

To perform the gel shift assay, protein nuclear extracts (1 μg) were incubated with 5-20 fmols of biotin labelled probe in 1x EMSA buffer (20 mM Tris HCl pH 7.5, 60 mM KCl, 1 mM EDTA, 1 mM DTT, 2 mM MgCl_2_, 25 μM ZnCl2, 0.1% Triton X-100, 5% glycerol, 200 μg/mL BSA) with 50 ng/μL poly (dI-dC) and 200 fmols of ssDNA oligo 5B1motF^38^ as unspecific competitors. Binding reaction was performed at room temperature for 20 min. If specific competition was assayed, incubation with an unlabelled probe was conducted for 20 min at room temperature prior to addition of the labelled probe. After binding, samples were run in a 4-5% acrylamide gel in 0.5x TBE buffer, blotted to a positively charged nylon membrane and developed using the Chemiluminescent Nucleic Acid Detection Module (Thermo Fisher) as recommended by the manufacturer.

### Purification of proteins bound to DNA fragments, proteomics and mass spectrometry analysis

Protein complexes bound to DNA fragments were isolated using magnetic Dynabeads MyOne Streptavidin C1 (Invitrogen) using a similar protocol as described before with some modifications^39^. Probes were generated with biotinylated oligos as for EMSA for 2.2 (ap2-g-3’PE) and 2.1 (control) regions (Fig. 1a). Biotin probes (100 pmoles) were then attached to the beads (50 μL) as recommended by the manufacturer, incubating DNA with the beads for 30 min at room temperature in binding and wash buffer (5 mM Tris HCl pH 7.5, 0.5 mM EDTA, 1M NaCl). Beads were then washed twice with the same buffer and twice with incomplete binding buffer (20 mM Tris HCl, pH 7.5, 60 mM KCl, 2 mM MgCl_2_, 0.1% IGEPAL CA-630). Protein nuclear extract (500 μL at 2 mg/mL) was then incubated with equilibrated beads (washed in incomplete binding buffer but with no probe) to perform a pre-clearing for 30 min at room temperature. Supernatant was then taken and mixed with 500 μL of 2x Binding buffer (20 mM Tris HCl, pH 7.5, 60 mM KCl, 2 mM MgCl2, 50 μM ZnCl2, 0.2% IGEPAL CA-630, 1 mM DTT, 2 mM EDTA, 1x protease inhibitor cocktail, 100 ng/μL poly(dI-dC)(dI-dC), 10% glycerol). The resulting mix (nuclear extract in binding buffer 1x) was split in two equal fragments (500 μL) and incubated for 30 min at room temperature with either 2.1 or ap2-g-3’PE probes attached to the beads. Next, seven washes were performed with 1 mL wash buffer (20 mM Tris HCl, pH 7.5, 250 mM KCl, 2 mM MgCl_2_, 25 μM ZnCl2, 0.1% IGEPAL CA-630, 1 mM DTT, 1x protease inhibitor cocktail, 5% glycerol), incubating in rotation for 5 min at room temperature each one. Three additional washes in 100 μL of ABC buffer (25 mM (NH_4_)HCO_3_) were performed keeping the beads in the magnet and with no incubation. Finally, beads were resuspended in 100 μL of ABC buffer and digested by adding 0.2 μg of trypsine/LysC (Promega) in 100 µL of ABC buffer for 1 hour at 37°C. Peptides were desalted and concentrated using homemade C18 StageTips before nano-liquid chromatography tandem mass spectrometry (LC-MS/MS) analysis. LC was performed with an RSLCnano system (Ultimate 3000, Thermo Scientific) coupled online to an Orbitrap Exploris 480 mass spectrometer (Thermo Scientific). Peptides were trapped on a C18 column (75 μm inner diameter × 2 cm; nanoViper Acclaim PepMap 100, Thermo Scientific) with buffer A (0.1% formic acid in DDW) at a flow rate of 3.0 µL/min over 4 min. Separation was performed on a 50 cm x 75 μm C18 column (nanoViper Acclaim PepMap RSLC, 2 μm, 100Å, Thermo Scientific) regulated to a temperature of 40°C with a linear gradient of 3% to 29% buffer B (100% MeCN in 0.1% formic acid) at a flow rate of 300 nL/min over 91 min. MS full scans were performed in the ultrahigh-field Orbitrap mass analyser in ranges m/z 375–1500 with a resolution of 120 000 at m/z 200. The top 20 most intense ions were subjected to Orbitrap for further fragmentation via high energy collision dissociation (HCD) activation and a resolution of 15 000 with the AGC target set to 100%. We selected ions with charge state from 2+ to 6+ for screening. Normalized collision energy (NCE) was set at 30 and the dynamic exclusion of 40s. For identification the data were searched against the UniProt *P. falciparum* (isolate 3D7: UP000001450) and the *Homo sapiens* (UP000005640) databases using Sequest^HF^ through proteome discoverer (version 2.2). Enzyme specificity was set to trypsin and a maximum of two missed cleavage sites were allowed. Oxidized methionine, N-terminal acetylation, and carbamidomethyl cysteine were set as variable modifications. Maximum allowed mass deviation was set to 10 ppm for monoisotopic precursor ions and 0.02 Da for MS/MS peaks. The resulting files were further processed using myProMS v3.9^40^. FDR calculation used Percolator and was set to 1% at the peptide level for the whole study. The label free quantification was performed by peptide Extracted Ion Chromatograms (XICs) computed with MassChroQ version 2^41^. For protein quantification, XICs from proteotypic peptides shared between compared conditions (TopN matching) were used, with missed cleavages allowed. Median and scale normalization was applied on the total signal of *P. falciparum* proteins to correct the XICs for each biological replicate (n=5). To estimate the significance of the change in protein abundance, a linear model (adjusted on peptides and biological replicates) based on two-tailed T-tests was performed and p-values were adjusted using Benjamini–Hochberg FDR procedure. *P. falciparum* proteins with at least three total peptides in all replicates, a 2-fold enrichment and an adjusted p-value < 0.05 were considered significantly enriched in sample comparisons.

## ACKNOWLEDGEMENTS

Authors wish to thank Dr. Flore Nardella for her technical support and advise on live imaging and Ms. Gretchen Diffendall for the critical revision of the manuscript. We also thank Mr. Florent Dingli for his support on proteomics mass spectrometry experiments.

This work was supported by a European Research Council advanced grant (grant PlasmoSilencing 670301) and the French Parasitology Consortium ParaFrap (grant ANR-11-LABX0024), awarded to A.S. C.C.-O is funded by the European Union’s Horizon 2020 research and innovation program under Marie Skłodowska-Curie grant agreement no. 741447.

## AUTHOR’S CONTRIBUTION

C.C.-O and A.S. conceived and designed the experiments and wrote the paper. C.C.-O performed the experiments and analysed the data. A.B. -S conducted RT-qPCR and contributed to ChIP-Seq experiments. A.C. carried out the gametocytogenesis conversion assays. C.S.-B developed the parasite mutant lines. D.L. and V.S. conducted the proteomics mass spectrometry and data analysis. All authors revised and approved the final manuscript.

## COMPETING INTERESTS

Authors declare no competing interests.

## DATA AVAILABILITY

Sequencing data from this study are available in the GEO (Gene Expression Omnibus) repository under accession number GSE145378. The proteomics mass spectrometry raw data have been deposited to the ProteomeXchange Consortium via the PRIDE^42^ partner repository with the dataset identifier PXD017930.

## Supplementary information

**Figure S1.**
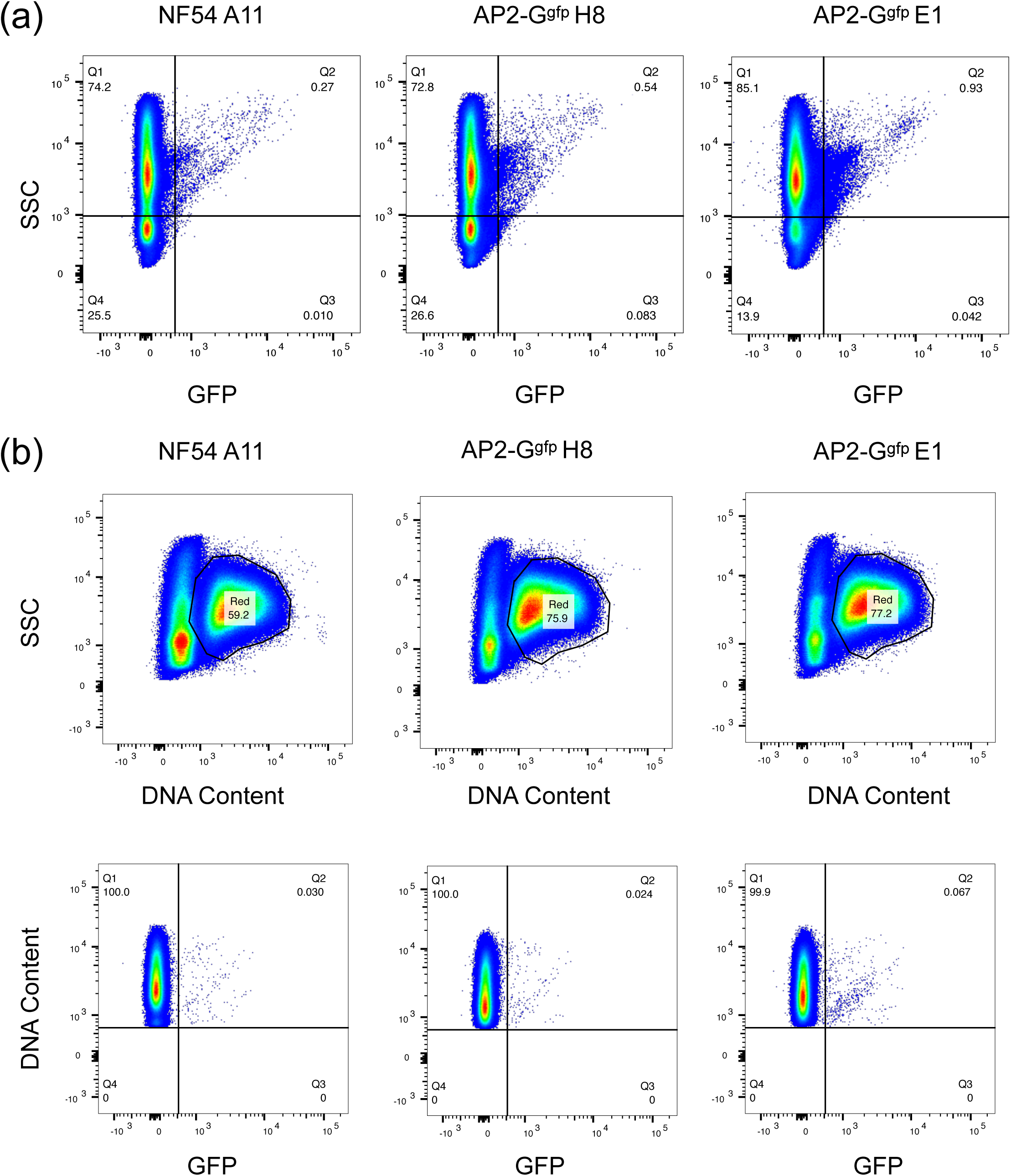
GFP detection by flow cytometry. (a) Whole population of infected RBCs from NF54 A11 and two clones of AP2-G_gfp_ line. (b) Same as (a) but gated parasites using Vibrant ruby staining.

**Table S1.**
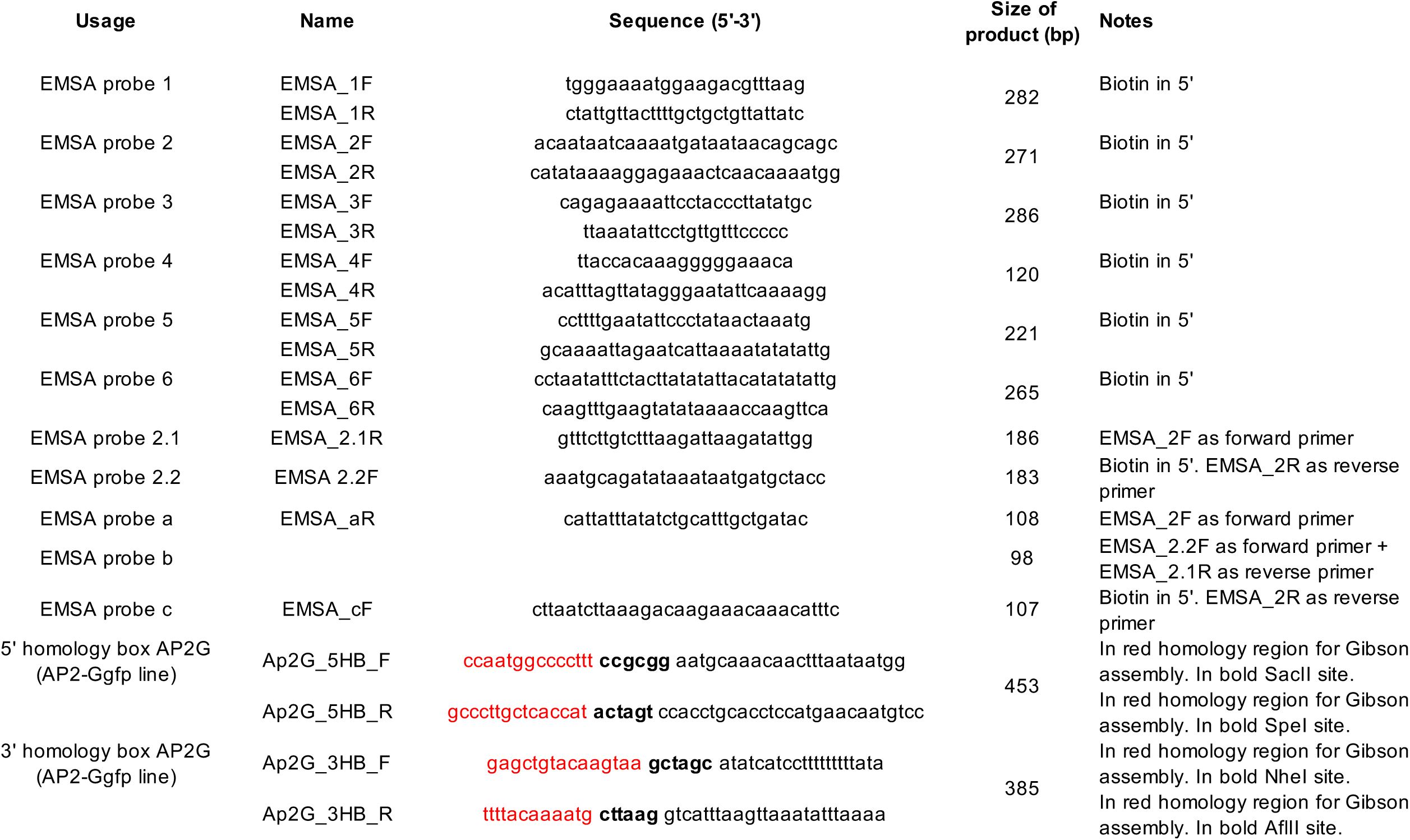
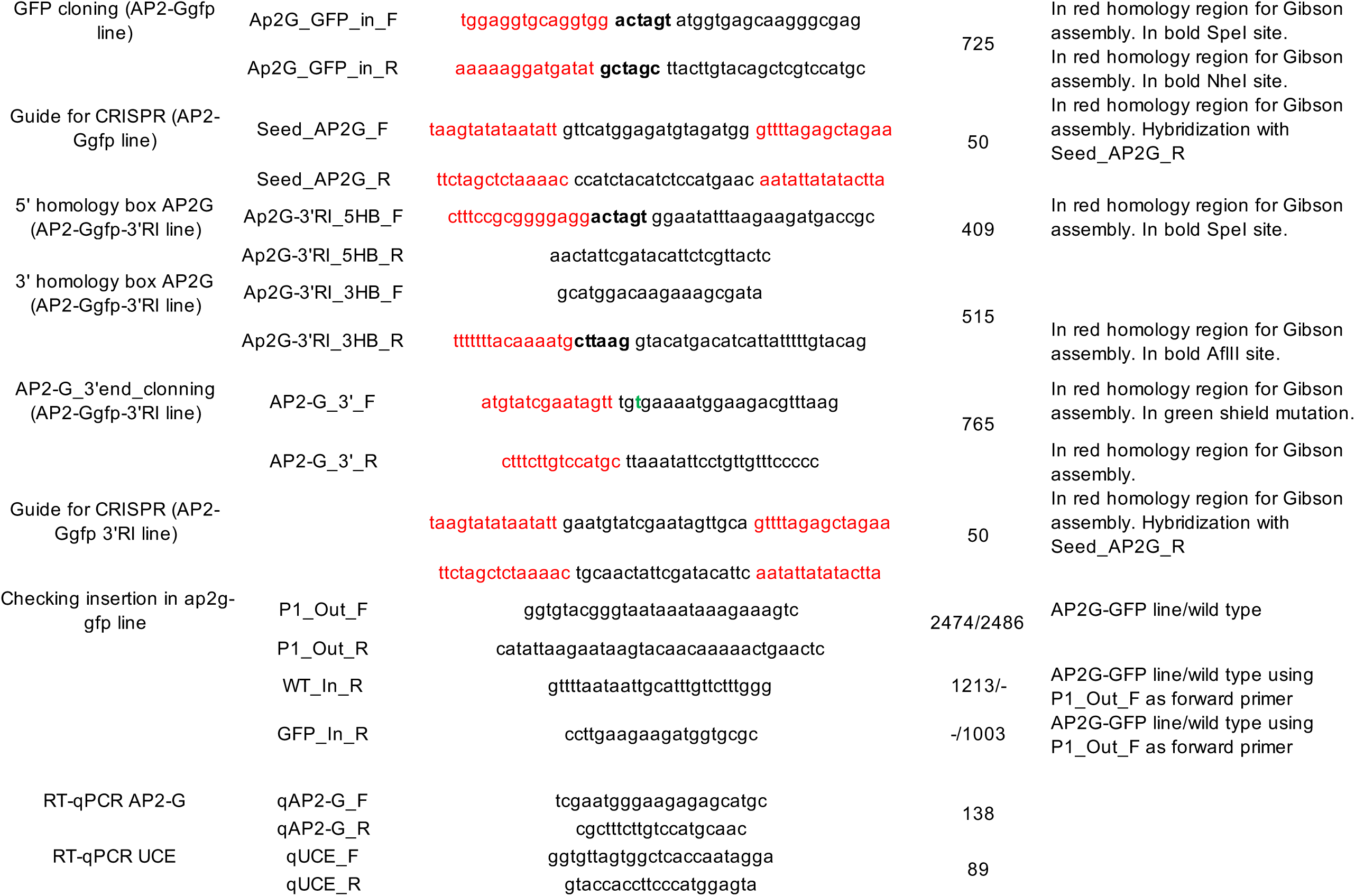
List of primers used in this work.

**Table S2.** Complete list of proteins found in the quantitative label-free mass spectrometry analysis.

| Gene | Sample/Control |  |  |  |  | Pept. used | Total Peptides | MW (kDa) | Description |
| --- | --- | --- | --- | --- | --- | --- | --- | --- | --- |
|  | Ratio | Log2 | Adj. p-value | CV % | Dist. pept. used |  |  |  |  |
| PF3D7_0523000 | 2.272258 | 1.184127 | 2.5E-119 | 2.993784 | 55 | 500 | 562 | 162.3 | Multidrug resistance protein 1 |
| PF3D7_0722200 | 3.850062 | 1.944882 | 3.96E-25 | 7.838067 | 20 | 212 | 220 | 87.9 | Rhoptry-associated leucine zipper-like protein 1 |
| PF3D7_0202000 | 2.317902 | 1.212819 | 1.57E-13 | 11.66072 | 21 | 210 | 235 | 71.3 | Knob-associated histidine-rich protein |
| PF3D7_0814200 | 3.239084 | 1.695586 | 1.46E-31 | 5.717741 | 16 | 130 | 153 | 27.3 | DNA/RNA-binding protein Alba 1 |
| PF3D7_1436300 | 3.190063 | 1.673585 | 9.37E-33 | 5.346129 | 14 | 119 | 128 | 112.4 | Translocon component PTEX150 |
| PF3D7_1004400 | 2.550159 | 1.350587 | 4.09E-14 | 9.885999 | 13 | 90 | 98 | 106.3 | RNA-binding protein, putative |
| PF3D7_1224300 | 2.173701 | 1.120153 | 1.1E-24 | 5.79835 | 10 | 81 | 104 | 97.2 | Polyadenylate-binding protein |
| PF3D7_1149000 | 2.788058 | 1.47926 | 4.64E-22 | 6.439867 | 9 | 79 | 86 | 689.3 | Antigen 332, DBL-like protein |
| PF3D7_1006200 | 2.270855 | 1.183235 | 4.23E-17 | 7.870744 | 7 | 72 | 81 | 12.0 | DNA/RNA-binding protein Alba 3 |
| PF3D7_0517300 | 2.239154 | 1.162954 | 1.95E-15 | 8.585299 | 9 | 71 | 82 | 34.6 | Serine/arginine-rich splicing factor 1 |
| PF3D7_0532400 | 5.489003 | 2.456544 | 6.14E-21 | 6.082335 | 7 | 64 | 71 | 61.1 | Lysine-rich membrane-associated PHISTb protein |
| PF3D7_0936800 | 2.046424 | 1.033105 | 2.73E-09 | 12.20996 | 5 | 54 | 55 | 45.5 | PRESAN domain-containing protein |
| PF3D7_1414300 | 2.183517 | 1.126654 | 1.79E-06 | 16.03374 | 6 | 53 | 66 | 25.2 | 60S ribosomal protein L10, putative |
| PF3D7_0827900 | 2.654137 | 1.408243 | 7.36E-15 | 7.675312 | 8 | 52 | 55 | 55.5 | Protein disulfide-isomerase |
| PF3D7_0920800 | 3.760727 | 1.911012 | 4.53E-17 | 5.845769 | 5 | 42 | 46 | 56.2 | Inosine-5'-monophosphate dehydrogenase |
| PF3D7_0501200 | 2.326734 | 1.218306 | 5.14E-07 | 14.13717 | 6 | 41 | 45 | 48.7 | Parasite-infected erythrocyte surface protein |
| PF3D7_1324900 | 2.001441 | 1.001039 | 2.8E-11 | 8.957916 | 5 | 39 | 43 | 34.1 | L-lactate dehydrogenase |
| PF3D7_0803700 | 2.298261 | 1.200543 | 5.11E-10 | 10.13789 | 4 | 38 | 40 | 51.6 | Tubulin gamma chain |
| PF3D7_0806800 | 1.99324 | 0.995115 | 1.18E-09 | 10.229 | 4 | 36 | 46 | 123.0 | V-type proton ATPase subunit a, putative |
| PF3D7_0904700 | 3.062787 | 1.614845 | 1.65E-11 | 6.953008 | 3 | 27 | 29 | 22.5 | Bacterial histone-like protein |
| PF3D7_0406200 | 2.051417 | 1.036621 | 3.65E-05 | 16.36308 | 3 | 26 | 29 | 16.6 | Sexual stage-specific protein |
| PF3D7_1006800 | 2.076046 | 1.053839 | 5.3E-07 | 9.783615 | 3 | 19 | 26 | 29.5 | Single-strand telomeric DNA-binding protein GBP2, putative |
| PF3D7_1321700 | 2.648584 | 1.405221 | 6.29E-06 | 12.06319 | 2 | 18 | 18 | 100.9 | Splicing factor 1 |
| PF3D7_0801000 | 6.574569 | 2.716896 | 7.73E-07 | 9.044315 | 3 | 17 | 21 | 147.1 | PRESAN domain-containing protein |
| PF3D7_0520400 | 2.351873 | 1.23381 | 0.002557 | 21.93887 | 2 | 17 | 19 | 35.7 | Single-stranded DNA-binding protein, putative |
| PF3D7_1115400 | 2.339434 | 1.226159 | 0.039746 | 34.72424 | 2 | 17 | 18 | 56.7 | Cysteine proteinase falcipain 3 |
| PF3D7_1113300 | 2.384632 | 1.253766 | 0.000744 | 18.32926 | 2 | 16 | 18 | 39.4 | UDP-galactose transporter, putative |
| PF3D7_0625700 | 2.377729 | 1.249584 | 0.001633 | 18.2784 | 2 | 13 | 14 | 12.0 | Uncharacterized protein |
| PF3D7_1310700 | 2.360856 | 1.23931 | 0.040534 | 32.87185 | 2 | 13 | 13 | 16.5 | RNA-binding protein, putative |
| PF3D7_1467100 | 36.06884 | 5.172681 | 2.57E-06 | 6.157749 | 1 | 10 | 10 | 59.4 | DNA-3-methyladenine glycosylase, putative |
| PF3D7_1353200 | 3.6563 | 1.870384 | 4.22E-07 | 3.797953 | 2 | 10 | 14 | 15.8 | Membrane associated histidine-rich protein 2 |
| PF3D7_1132300 | 3.230581 | 1.691794 | 0.001465 | 15.917 | 1 | 10 | 10 | 31.7 | Nucleic acid binding protein, putative |
| PF3D7_0415900 | 3.155587 | 1.657908 | 0.000518 | 13.41018 | 1 | 10 | 10 | 24.1 | Ribosomal protein L15 |
| PF3D7_0416900 | 1000 | 1000 |  |  | 2 | 9 | 9 | 289.7 | Uncharacterized protein |
| PF3D7_1138800 | 1000 | 1000 |  |  | 2 | 9 | 9 | 222.1 | WD repeat-containing protein, putative |
| PF3D7_1323700 | 3.670588 | 1.876011 | 0.001613 | 14.83887 | 1 | 9 | 10 | 34.0 | Glideosome associated protein with multiple membrane spans 1 |
| PF3D7_1142600 | 2.262074 | 1.177646 | 9.48E-05 | 9.030268 | 1 | 9 | 10 | 16.3 | 60S ribosomal protein L35ae, putative |
| PF3D7_0624400 | 2.124958 | 1.087434 | 0.000141 | 9.663783 | 1 | 9 | 10 | 73.1 | Uncharacterized protein |
| PF3D7_1143300 | 2.229411 | 1.156662 | 2.8E-05 | 5.945781 | 1 | 8 | 10 | 39.0 | DNA-directed RNA polymerases I and III subunit RPAC1, putative |
| PF3D7_0316600 | 6.123615 | 2.614384 | 0.00147 | 10.72317 | 1 | 7 | 10 | 34.5 | Formate-nitrite transporter |
| PF3D7_1121100 | 4.059666 | 2.021361 | 0.011383 | 18.02303 | 1 | 7 | 8 | 184.9 | Uncharacterized protein |
| PF3D7_0310600 | 2.09436 | 1.06651 | 0.00822 | 13.58612 | 2 | 7 | 9 | 28.0 | Eukaryotic translation initiation factor 3 subunit K |
| PF3D7_1148900 | 3.909436 | 1.966961 | 0.046695 | 24.33332 | 1 | 6 | 8 | 10.3 | Uncharacterized protein |
| PF3D7_1109900 | 2.632921 | 1.396664 | 0.048485 | 24.69078 | 1 | 6 | 8 | 12.8 | 60S ribosomal protein L36 |
| PF3D7_0515700 | 1000 | 1000 |  |  | 2 | 5 | 5 | 51.8 | Glideosome-associated protein 40, putative |
| PF3D7_1415400 | 1000 | 1000 |  |  | 1 | 5 | 5 | 178.5 | Btz domain-containing protein |
| PF3D7_1370300 | 1000 | 1000 |  |  | 1 | 4 | 4 | 28.9 | Membrane associated histidine-rich protein 1 |
| PF3D7_0614200 | 1000 | 1000 |  |  | 1 | 4 | 4 | 107.9 | Cytosolic Fe-S cluster assembly factor NAR1, putative |
| PF3D7_1222700 | 1000 | 1000 |  |  | 1 | 4 | 4 | 23.6 | Glideosome-associated protein 45 |
| PF3D7_1364800 | 1000 | 1000 |  |  | 1 | 4 | 4 | 24.2 | DNA-directed RNA polymerases I, II, and III subunit RPABC1, putative |
| PF3D7_1342600 | 1000 | 1000 |  |  | 1 | 4 | 4 | 92.3 | Myosin-A |
| PF3D7_1102500 | 1000 | 1000 |  |  | 1 | 4 | 4 | 73.9 | PRESAN domain-containing protein |
| PF3D7_1411100 | 1000 | 1000 |  |  | 1 | 4 | 4 | 44.2 | Uncharacterized protein |
| PF3D7_1234000 | 1000 | 1000 |  |  | 1 | 3 | 3 | 50.7 | Uncharacterized protein |
| PF3D7_1142300 | 1000 | 1000 |  |  | 1 | 3 | 3 | 216.0 | Uncharacterized protein |
| PF3D7_1022300 | 1000 | 1000 |  |  | 1 | 3 | 3 | 37.2 | ZIP domain-containing protein, putative |
| PF3D7_1465000 | 1000 | 1000 |  |  | 1 | 3 | 3 | 33.3 | Cytochrome c oxidase subunit ApiCOX25, putative |
| PF3D7_1440700 | 1000 | 1000 |  |  | 1 | 3 | 3 | 87.2 | AP-3 complex subunit mu, putative |

## Notes

### Competing Interest Statement

The authors have declared no competing interest.

